# Walk the Plank! Using mobile EEG to investigate emotional lateralization of immersive fear in virtual reality

**DOI:** 10.1101/2022.08.30.505699

**Authors:** Yasmin El Basbasse, Julian Packheiser, Jutta Peterburs, Christopher Maymon, Onur Güntürkün, Gina Grimshaw, Sebastian Ocklenburg

## Abstract

Most studies on emotion processing rely on the presentation of emotional images or films. However, this methodology lacks ecological validity, limiting the extent to which findings can generalize to emotion processing in the wild. More realistic paradigms using Virtual Reality (VR) may be better suited to investigate authentic emotional states and their neuronal correlates. This preregistered study examines the neuronal underpinnings of naturalistic fear, measured using mobile electroencephalography (EEG). Seventy-five healthy participants entered a simulation in which they walked across a virtual plank which extended from the side of a skyscraper – either 80 stories up (the negative condition) or at street level (the neutral condition). Subjective ratings showed that the negative condition induced feelings of fear and presence. Following the VR experience, subjects passively viewed negative and neutral images from the International Affective Picture system (IAPS) outside of VR. We compared frontal alpha asymmetry between the plank and IAPS task and across valence of the conditions. Asymmetry indices (AI) in the plank task revealed greater right-hemispheric lateralization during the negative VR condition, relative to the neutral VR condition and to IAPS viewing. Within the IAPS task, no significant asymmetries were detected, though AIs in the VR task and in the IAPS task were negatively correlated suggesting that stronger right-hemispheric activation in the VR task was associated with stronger left-hemispheric activation during the IAPS task. In summary, our findings indicate that immersive technologies such as VR can advance emotion research by providing more ecologically valid ways to induce emotion.

## 2. Introduction

Since early hints from lesion research [1–3], many studies have supported the notion that emotion processing in the brain is lateralized [4]. Despite the consensus that emotions are processed asymmetrically in the brain, an exhaustive theory explaining the underlying mechanisms is yet to be found. Rather, there are several theories each holding strong empirical support for different aspects of emotional experience but also showing considerable inconsistencies. For example, the two most-established theories about emotion lateralization are the right hemisphere and the valence hypothesis. The former postulates that the right hemisphere is responsible for emotional experiences irrespective of valence [5–10], whereas the latter differentiates between the valence of the emotion with positive emotions being processed in the left hemisphere and negative emotions being processed in the right hemisphere [11–15]. To this day, studies gather evidence for both theories even though the core predictions are not complementary [16].

One reason for these mixed findings could be the lack of ecological validity in the operationalization of emotions. In laboratory settings, the most popular tools to elicit emotions experimentally are images (e.g., IAPS [17]) or emotional films. Adding moving pictures should increase immersion and saliency of the stimuli thereby affecting emotions more strongly than static images [18]. In general, both images and films were shown to elicit behavioral and physiological emotional reactions (images [19,20]; films [21,22]). However, their repetitive presentation in block designs conflicts with an ecologically valid understanding in which emotions are characterized by their spontaneous and phasic nature. Moreover, the mere perception of emotional stimulus material does not reflect the multi-faceted nature of emotional experience. In real life, perceiving emotional content quickly translates into and prepares the body for action like the fight-or-flight response [23]. These crucial components of emotion are neglected by studies focusing solely on emotion perception.

One potential solution to this problem is the use of Virtual Reality (VR). VR is already widely used for treatment in clinical populations, for example, for exposure therapy in fear of heights [24], social phobia [25,26], or eating disorders [27]. In healthy participants, VR is often used for educational purposes [28–30] or to investigate spatial navigation [31–33]. One of the major advantages of VR is the possibility to simulate real-life situations realistically while minimising external influences. Thereby, VR bridges experimental control and ecological validity. Because the environment responds to participants’ movements, VR offers a higher degree of immersion into the experimental paradigm compared to two-dimensional content, making it effective in eliciting “authentic” emotions [34–36]. Fear induction, particularly, can benefit from more realistic methods like VR. Although fear has been successfully elicited in prior experiments as evident in self-report and bodily reactions like elevated heart rate or skin conductance [19,20], the selection of suitable (but also ethical) stimulus material or situations can be challenging. For example, Barke, Stahl, and Kröner-Herwig [37] found that some IAPS images selected by experts to evoke fear (especially images depicting human threat) actually provoked anger in the participants. Moreover, fear evoked by pictures does not have the intensity of genuine threat. By using interactive simulations resembling fearful situations in real life, VR could offer a more valid approach to experimental fear induction. This in turn could further widen the scope of emotion research from mere perception to perception-to-action.

In order to investigate the neuronal underpinnings of emotional experience, VR can be combined with mobile EEG systems [38,39]. These systems possess acceleration sensors specifically designed to take movement into account which can then be integrated in data analysis. Thereby, a major drawback of stationary EEG studies for emotion research can be addressed. In a conventional EEG system, the electrodes are connected to the amplifier via cables so that it is usually necessary that participants lay their head on a chin rest to avoid movements as much as possible. This, however, severely limits the spectrum of paradigms suitable for EEG experiments [40] particularly affecting emotion research. Natural spontaneous reactions such as frowning, smiling, wincing, or body movements which are indicative of experiencing real emotions become disadvantageous for later data analysis and are sought to be suppressed. In contrast, the largely wireless signal transmission of mobile EEG systems allows for free movement without notable interference with the neuronal signal [41–43], meaning that researchers can take advantage of both the heightened perceptual experience in VR, and the freedom of movement it affords.

In the present preregistered study, we aim to investigate the neuronal correlates of experimentally induced fear in two paradigms with varying levels of ecological validity. The first task comprises a naturalistic VR setting with high ecological validity. Here, participants balance either on a virtual plank on a tall skyscraper (negative condition) to induce fear of heights, or on the ground (neutral condition). The second task comprises a typical fear induction paradigm with low ecological validity during which the participants are presented with fearful or neutral IAPS images. As both major theories in emotional lateralization (right hemisphere and valence hypothesis) predict a stronger right-hemispheric activation in situations of negative valence, we predict that stronger rightward activation in both tasks. Since stronger experiences of emotions are furthermore associated with higher levels of asymmetry for negative emotions [44], the more realistic fear evoked by the VR task could also be reflected in a more lateralized rightward activation pattern compared to the non-realistic IAPS task.

Based on previous research on emotion processing, hemispheric asymmetries are quantified using alpha power. Alpha power can be derived from the EEG recording and is inversely correlated with cognitive activity. Higher alpha power on a given electrode site indicates relatively less activity in that region [45]. Since emotional content requires attentional and cognitive resources, alpha power is expected to decrease as a function of emotional intensity. To deduce hemispheric asymmetries, alpha power from homologous regions in both hemispheres can be compared using the asymmetry index, particularly for frontal electrode pairs F3/F4 [46,47] and F7/8 [41]. Complementing the neurophysiological measures, fear ratings are acquired during the VR task.

We predict that alpha at F3/F4 and F7/F8 will show greater relative left-sided power in the negative VR condition compared to the neutral VR condition. This would reflect right-hemispheric processing of negative emotion as predicted both by the right hemisphere and valence hypotheses. Similarly, greater left than right alpha power at the electrode pairs F3/F4 and F7/F8 is predicted for the negative condition of the IAPS task compared to the neutral condition. It will be explored whether alpha asymmetries in the VR task are more pronounced overall than in the IAPS task as a response to the more realistic fear exposure. If alpha asymmetry supports fear experience, we further predict that the self-reported levels of fear should positively correlate with higher left-sided alpha power indicative of stronger right-hemispheric activation. Finally, we predict that alpha power asymmetries should show higher correlations during the neutral conditions of both tasks due to similar (non)-emotional processing, relative to the negative conditions of both tasks, given the expected difference in negative emotions due to differences in ecological validity. All hypotheses and analyses were pre-registered at Open Science Framework (https://osf.io/uv26f).

## 3. Materials and Methods

### 3. 1 Participants

We aimed to collect data from 80 participants prior to data analysis. A power analysis of a within-subject repeated-measures ANOVA with three measurements and four groups revealed that in this sample size it is possible to detect small to medium sized effects (Cohen’s f = 0.16) at 80% power. Ninety-three potential participants filled out an online survey to take part in the experiment. Exclusion criteria were age (below 18 and above 35 years), diagnosis of mental/neurodevelopmental disorders or neurological diseases (e.g., migraine, epilepsy), and extreme fear of heights. One person did not fit the age criterion and thus could not participate, and one person was excluded a posteriori due to being older than 35 years. There were four cases in which mental disorders led to an exclusion from the study, namely one person whose score on the Beck Depression Inventory (BDI [48]) was above the cut-off value (51; a value ≥ 30 indicates severe depression) and three persons reporting bipolar disorder, panic disorder, and depression with addiction. These measures were taken due to empirical evidence revealing altered hemispheric asymmetries in clinical samples compared to non-clinical samples [49,50]. For the same reason, high values in the Acrophobia Questionnaire (ACRO/AVOI [51]; German version [52]) were considered an exclusion criterion which affected one potential participant. After applying the exclusion criteria, the sample comprised 87 healthy participants (54 women, 33 men) with normal or corrected-to-normal vision. In a second step, participants were excluded due to issues during data acquisition or data analysis. In one case, the experiment had to be cancelled during the VR condition, because the subject reported sudden anxiety symptoms and could not complete the task. In two cases, technical difficulties with the IAPS paradigm occurred leading to exclusion from data analysis. EEG pre-processing revealed missing triggers in four and incorrect markers in two EEG recordings which also led to the exclusion of these files from further analysis. These technical issues were likely caused by signal loss in the connection of the mobile EEG system to the recording laptop. One person was excluded from analysis because the screening questionnaire was not filled out. Thus, the final sample used for statistical analysis comprised 75 subjects (45 women, 30 men) with a mean age of 24 (SD = 3.61). A sensitivity analysis showed that this sample retained the ability to detect small to medium effects (Cohen’s d = 0.34) at 80% power.

The study was accepted by the ethics committee of the psychological faculty of the Ruhr-University Bochum. All participants gave their written informed consent at the beginning of the experiment. The experiment was conducted in accordance with the Declaration of Helsinki.

### 3. 2 Stimulus material

#### 3. 2. 1 IAPS paradigm

Two sets of stimuli were chosen from the IAPS database (https://csea.phhp.ufl.edu/index.html), one consisting of 50 neutral images and one with 50 negative images. Negative images were picked according to their arousal and valence ratings determined by previous validation studies [17,53]. In the negative sample, only images with an arousal rating over 4.3 (M = 6.33; SD = .67) and a valence rating under 2.54 (M = 2; SD = .41) were included to ensure that the pictures reliably induce negative affect. A list of the selected slides and ratings can be found in the Supplementary material D. Negative stimuli mostly contained pictures displaying mutilation, injuries, and violent scenes. The neutral stimulus set was composed of pictures showing mainly household objects, nature, and different patterns. The IAPS conditions were programmed using the software Presentation.

#### 3. 2. 2 Virtual Reality paradigm

The VR simulation was presented using an HTC Vive HMD with corresponding hand controllers (https://www.vive.com/us/). The VR environment was created in the lab using the Unity game engine (https://unity.com/). The simulation was projected to the HMD using the SteamVR [54] runtime application. The VR set up included two base stations framing the virtual room, the HMD, two controllers, and one laptop with an external number pad to choose VR conditions. Apart from that, there was one laptop to display the IAPS task. One additional laptop was needed for EEG signal recording.

### 3. 3 Procedure

Participants were tested in a designated test room in the Biopsychology department at Ruhr University Bochum. The duration of the experiment was approximately two hours. Due to the ongoing Covid-19 pandemic at the time of the inquiry, participants were obliged to undergo a Covid-19 self-test and to wear a mask ensuring minimal infection risk for both participant and experimenter. The participants were then asked to fill out the written informed consent, a Covid-19 symptom questionnaire, and an attendance list to track contacts in the event of a positive Covid-19 case. The LiveAmplifier was placed in a small pocket at the back of the EEG cap. After a secure wireless connection between LiveAmplifier and recording laptop was established, the EEG cap was prepared with electrode gel to allow signal detection and transmission. To connect the LiveAmplifier with the Sensor & Trigger Extension box, participants wore a belt bag containing the latter and a power bank to guarantee power supply of the equipment. The first EEG recording captured resting state EEG while sitting. Participants were asked to close their eyes for five minutes and relax without concentrating on anything particular. During this time, the experimenter left the room and turned off the lights to avoid distraction. After five minutes elapsed, subjects filled out the first PANAS and the STAI-S questionnaire.

To prepare for the VR task, a wooden plank (width: 19 cm; depth: 4 cm; length: 151 cm) was placed in the center of the room. The plank was manipulated so that it was slightly unstable, wobbling to the left and right at each step. Before entering VR, participants had the chance to walk on the plank once to eliminate novelty effects. Thereafter, participants were introduced to the two affective ratings (presence in VR and subjective fear, see 3.4.2 for details) inquired during the VR task. The experimenter asked for the first rating to check if the ratings were understood correctly. Subsequently, the HMD was fitted to the EEG cap and the participant was guided to the starting point of the VR task. The order of the negative and neutral condition was counterbalanced. In both conditions, the participants first found themselves standing in an open elevator on ground level looking at a city scene. After ensuring that the simulation started properly, the participants were handed two controllers. To enhance presence in the VR, they were told to make themselves familiar with them by operating their virtual hands and fingers. Apart from the virtual environment, the experimental condition and control condition followed an identical protocol.

Once inside the virtual environment, participants were asked to step outside the elevator slowly moving toward the street. The participants then turned around and re-entered the elevator. Participants then used their right hand controller to press a red button on the right side of the elevator door, causing the elevator doors to close and the elevator to move upwards. In the neutral condition, the elevator moved upwards only for a short period before returning to the street level again. In the negative condition, the elevator continued to the top of the building. Opening of the elevator doors marked the beginning of a one minute segment in which the participant was told to look around without walking or talking (see Figure 1A (neutral condition) and 1C (negative condition) for the participant perspective). The manual EEG trigger was pressed to set accurate timepoints for subsequent analyses. After one minute, the experimenter asked for the first rating (‘elevator’ rating). Next, the participants took one step onto the plank and stopped there, initiating the next one-minute-interval of looking around (see Figure 1B (neutral condition) for the participant perspective). Having completed one minute, participants were asked to give their second rating (‘start of plank’ rating). Afterwards, participants were instructed to walk further along the plank until the end of the plank was reached (see Figure 1D (negative condition) for the participant perspective). Stopping there marked the beginning of the third one-minute-long recording interval of the EEG. After one minute, participants gave their rating again (‘end of plank’ rating). Subjects could then walk back on the plank and re-enter the elevator thereby finishing the respective condition.

**Figure 1.**
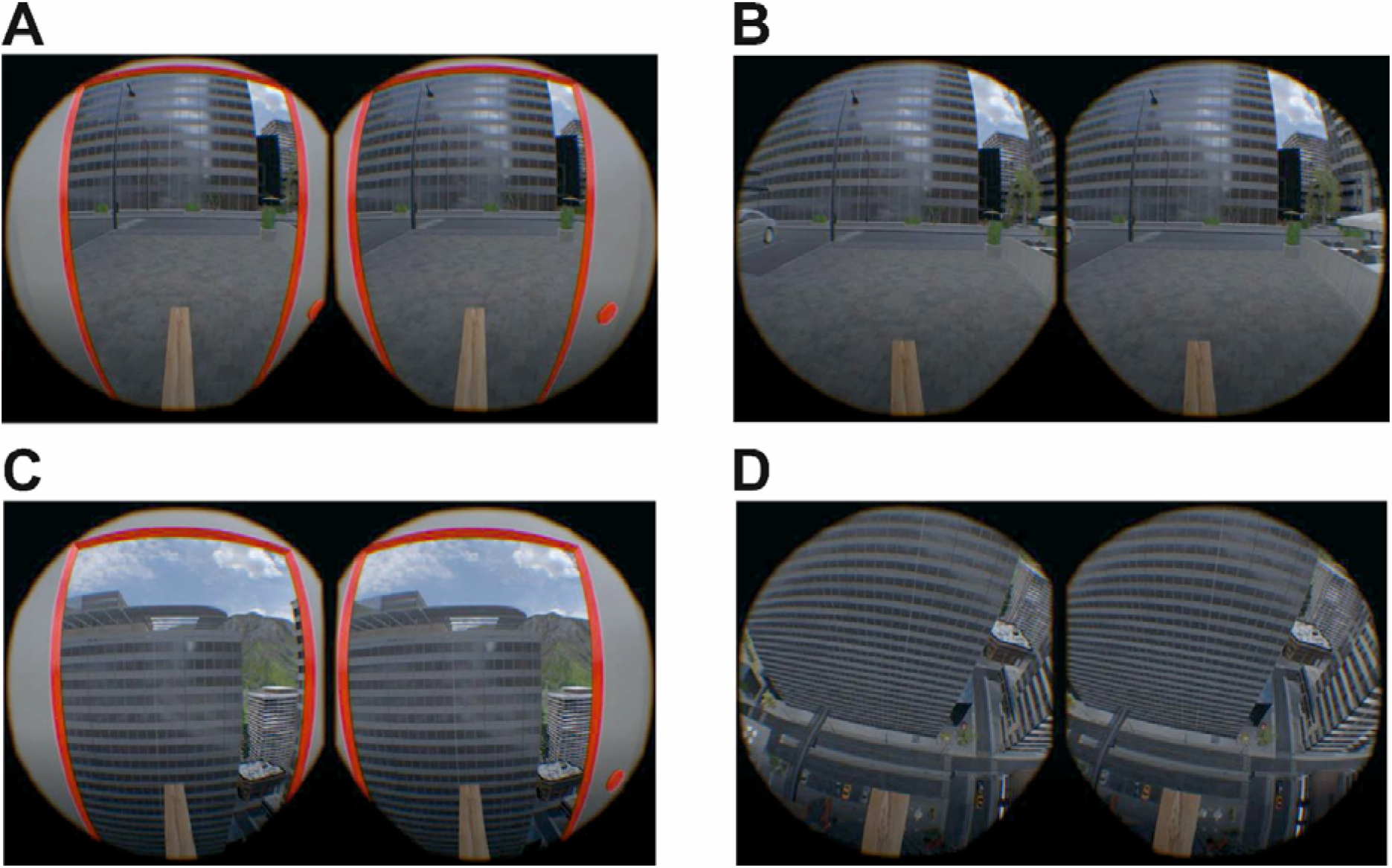
Example views from the participants’ perspective. **A)** Street view from inside the elevator in the neutral condition. **B)** Street view in the start of plank recording segment in the neutral condition. **C)** View from the building top from inside the elevator in the negative condition. **D)** Downwards view from the end of plank recording segment in the negative condition.

It needs to be noted that the experimental paradigm employed in the study deviated from the outlined design in the pre-registration. In the pre-registration, six time points for EEG recordings were planned (at street level, bottom of elevator, top of elevator, on plank, end of plank, turning back). These were however reduced to three as two of these conditions would have had complete overlap between the experimental and control condition (street level and bottom of elevator). The turning back condition was removed to avoid a condition with considerable movement artifacts as the participant would have manoeuvred on the plank.

Having completed the VR task, the HMD was loosened from the EEG cap. Participants could take a seat to fill out the second PANAS questionnaire and to do the second part of the experiment. The manual trigger was unplugged from the trigger box and the latter was connected to the presentation laptop so that automatic triggers embedded in the IAPS presentation could be transmitted to the recording laptop. The proper IAPS condition was prepared following the counterbalanced order, starting either with neutral or with negative pictures. The participants were briefed to observe the pictures attentively and thoroughly without looking around the room. The presentation started by pressing the spacebar on the presentation laptop. Simultaneously, the experimenter ran the new EEG file. During the presentation, the experimenter left the room and turned the lights off to avoid distraction. Each picture was displayed for 5 s with an interstimulus interval of 2 s, leading to a total trial duration of 7 min for each condition. Directly following the first set of pictures, the second set was started, and the procedure described above was repeated. When the second set was finished, participants were instructed to fill out the post-experiment survey via Qualtrics. Subsequently, the EEG equipment (i.e., belt bag, EEG cap) was removed from the participants, and they were compensated either with 20€ or course credit. The timeline of the experiment is illustrated in Figure 2.

**Figure 2.**
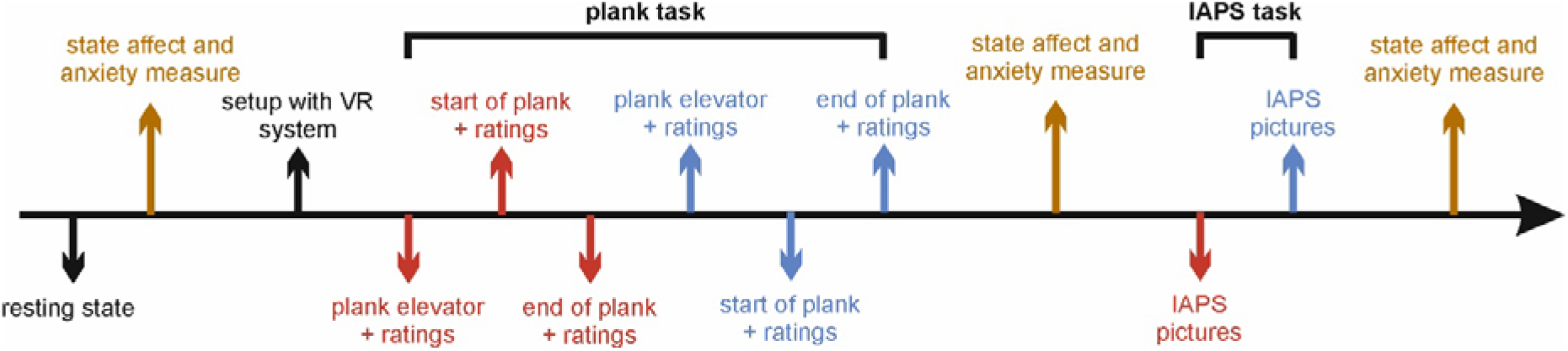
Timeline of the experimental procedure. Before the experiment, participants first went through a resting state before providing information about state affect and anxiety. After being set up with the VR equipment, the plank task was conducted. Note that the red and blue color illustrate the negative and neutral conditions which were counterbalanced across participants. Ratings include the presence as well as subjective fear rating. After completing the plank task, another state affect and anxiety measure was taken. The following IAPS task also comprised a negative and neutral condition (counterbalanced across participants). Finally, state affect and anxiety were measured before the end of the experiment.

### 3. 4 Behavioral Measures

#### 3. 4. 1 Questionnaires

Potential participants were sent a link to an online screening on the platform Qualtrics (https://www.qualtrics.com/) for relevant exclusion criteria. The screening contained demographic information, exclusion criteria, and a Covid-19 risk and symptom screening (see Supplementary material A). In terms of demographic information, age, gender, vision, spoken languages, educational achievement, and mental and physical health were inquired. Furthermore, various questionnaires relevant for later analyses were included. The State-Trait Anxiety Inventory-Trait (STAI-T [55]) measures self-reported trait anxiety with 20 items. On a 4-point Likert scale (from “not at all” to “very much so”), participants determine to what extent they generally agree with different anxiety-related statements (e.g., “I feel rested.”) resulting in a trait anxiety score from 20-80. The Acrophobia Questionnaire (ACRO/AVOI [52]) assesses the self-reported extent of fear (ACRO) and avoidance (AVOI) of 20 specifically height-related situations (e.g., standing on the edge of a railway track, riding a Ferris wheel, or standing on a balcony on the 10th floor). For each situation, participants report how anxious they would feel given that situation and to what extent they would avoid it on a 7-point Likert scale (from “not at all anxious/would not avoid doing it” to “extremely anxious/would not do it under any circumstances”). Consequently, a mean score can be calculated each for fear and avoidance of heights. It is important to note that the ACRO/AVOI was not designed to differentiate between clinically relevant acrophobia and non-clinical unease of heights. After completing the screening, subjects were contacted via e-mail to arrange an appointment for the experiment.

On site, participants filled out the state version of the STAI (STAI-S [55]). The STAI-S resembles the trait version (STAI-T) regarding the items but assesses state anxiety in that given moment. This way, baseline anxiety before experimental fear induction was documented. To assess overall affective states, the Positive and Negative Affect Schedules (PANAS, German version [56]; English version [57]) was given to the participants. The PANAS comprises of 20 adjectives describing different feelings and moods (e.g., strong, worried, or proud). On a 5-point Likert scale from “not at all” to “extremely”, participants tick the box which best describes their feelings in that moment. All questionnaires can be found in Supplementary material B.

Finally, the Simulator Sickness Questionnaire [58] and the iGroup Presence Questionnaire [59] (see Supplementary material C) were filled out by the participants. These questionnaires evaluate to what extent participants felt immersed and affected by the VR simulation. The Simulator Sickness Questionnaire evaluates on a 4-point Likert scale (from “not at all” to “strongly”) to what degree participants experienced 16 symptoms characteristic for simulations in VR, e.g., blurred vision, dizziness (with eyes open or closed), or general discomfort. The Sense of Presence Inventory assesses how present participants felt in the virtual environment and if a sense of “being there” [59] could be induced. To do so, participants were asked, for example, how aware they were of the real world while being in the virtual world.

#### 3. 4. 2 Affective ratings

Two kinds of ratings were carried out addressing subjective fear and presence. Ratings were given within a range from 1 (not fearful/present) to 10 (extremely fearful/present) at three recording segments (elevator, start of plank, end of plank). Before entering the VR simulation, a test fear rating was implemented to check whether participants had understood the ratings. This rating was not included in the statistical analysis.

### 3. 3 Physiological measures

Brain activation was measured with a mobile EEG system (LiveAmp 32, Brain Products GmbH, Gilching, Germany). This system consists of 32 electrodes arranged in line with the international 10-20 system [60]. The position of the Fpz electrode was set as the ground electrode, and FCz served as the reference electrode. A wireless amplifier amplified the recorded activation transmitting it to the recording software (Brain Vision Recorder) on a laptop with a sampling rate of 1000 Hz. The impedance cut-off was set to <10 KΩ for an adequate EEG signal. Three acceleration sensors implemented in the wireless amplifier measured head and body movements along the X, Y, and Z axes to enable subsequent movement correction in the recordings. Several EEG caps in different sizes were adjusted for VR-compatible use by MES Forschungssysteme GmbH (https://mes.gmbh/). To this end, five Velcro straps were sewed onto each cap attaching the head-mounted display (HMD) without interfering with the electrodes.

### 3. 6 EEG Pre-processing

For data preprocessing, the program BrainVision Analyzer (Brain Products GmbH, Gilching, Germany) was used. First, the sampling rate was adjusted from 1000Hz to 250Hz with a sampling interval of 4000μS to shorten processing time. A 0.5Hz high-pass filter (zero phase shift Butterworth filter, time constant 0.3183099) was applied to avoid interference from non-physiological high frequency sources. To exclude noise from electronic devices in the experimental setting, a 50Hz notch filter was used. First, electrodes that showed notably low (i.e., flat lines) or high (i.e., unusual spikes) variance were removed from the recording. Then, these channels were interpolated using the surrounding electrodes. Raw data was inspected manually, detecting and removing artifacts caused by coughing or excessive movement. An infomax independent component analysis (ICA) extracted pulse and eye artifacts (i.e., blinking and horizontal eye movement). The reference electrode FCz was interpolated. The data from the plank task was then segmented according to the three markers (S1: elevator; S2: start of plank; S3: end of plank) set with the manual trigger. Segment length of the first two markers varied since each following marker constituted the end of the prior interval (e.g., S2 marked the end of the segment starting with S1). The segment period for the third marker was 60000ms. Overlap of segments was not allowed. The average of all channels was then set as the new reference (Avg). A second filter was applied with a low cut-off at 1Hz and a high cut-off at 45Hz. Segments with a length of 1s were created via segmentation. A Fast-Fourier Transformation (FFT; Hanning window of 10%) was applied to extract alpha oscillations (8-12Hz) from the signal. Lastly, the average power was calculated for each recording segment in the plank conditions. In the IAPS task, the preprocessing steps were the same as for the plank task except that there were no segments following marker positions. Segments of 1s were created for the whole length of the paradigm regardless of stimulus markers. Consequently, only one average was calculated for the whole IAPS condition for later power analysis.

For movements, no initial IIR filter was applied to enable analysis of the raw data. All electrodes except for the three acceleration sensors were disabled. In the plank task, data from the acceleration sensors was segmented following the same logic based on the marker positions of S1, S2, and S3 described above. In a second step, segments with a length of 1s were defined. For each recording segment, an average was calculated. Data from each acceleration sensor (X-axis, Y-axis, Z-axis) was exported separately for the recording segments. The same procedure was applied to the movements in the IAPS task. There was again one average for the whole condition from which the data for the acceleration sensors was exported separately. To investigate hemispheric asymmetries in the experiment, the asymmetry index was calculated for the electrode pairs F3/4 and F7/8. The asymmetry index is obtained by subtracting the natural logarithm of the frequency band power of a specific left-hemispheric electrode site from the natural logarithm of the frequency band power of its right-hemispheric homologue [61]. The formula is as follows:

AI = ln(power right)-ln(power left)

A positive alpha index indicates greater relative left than right activation and a negative alpha index indicates greater relative right than left activation as alpha is inversely correlated with brain activity [45].

### 3. 7 Statistical analyses

#### 3. 7. 1 Pre-registered analyses

##### Hypotheses 1 and 2

A one-group repeated-measures design was used with the within-subject factors condition (negative, neutral), validity (plank, IAPS), and electrode pair (F3/F4, F7/F8). Therefore, a 2 × 2 × 2 repeated measures ANOVA with the described factors was calculated. For the plank task, a repeated-measures 2 × 2 × 3 ANOVA with the within-subject factors valence (neutral, negative), electrode (F3/4, F7/8), and recording segment (‘elevator’, ‘start of plank’, ‘end of plank’) was applied. Since the pre-registration had six instead of three recording segment levels, the pre-registration originally planned a 2 × 2 × 6 ANOVA. Since the experimental protocol was changed prior to data collection, the analysis design was adapted accordingly. Repeated-measures ANOVAs were Greenhouse-Geisser corrected if sphericity was violated. Significant main and interaction effects were corrected post hoc using the Bonferroni method.

##### Hypotheses 3 and 4

Self-reported fear ratings were correlated with the AIs in the corresponding VR condition and recording segment. AIs of the neutral and the negative IAPS condition were correlated with the AIs of the neutral and negative plank condition.

#### 3. 7. 2 Exploratory analyses

##### 3. 7. 2. 1 Self-report data

For the VR task, a 2 × 3 repeated-measures ANOVA of fear ratings with the within-subject factors valence (neutral, emotional) and recording segment (‘elevator’, ‘start of plank’, ‘end of plank’) was calculated. Fear ratings were correlated with the ACRO/AVOI scale, the trait and state version of the STAI, the SSQ, and with presence ratings. Presence ratings were correlated with the IPQ from the screening survey. A repeated-measures 3 × 2 ANOVA of the three PANAS questionnaires with the within-subject factors time (baseline, post-VR, post-IAPS) and affect (positive, negative) was conducted.

##### 3. 7. 2. 2 Movement signals

For every condition (neutral, negative) and electrode pair (F3/4, F7/8), a multiple linear regression with the AI as dependent variable and the individual acceleration sensor signals (X, Y, and Z) as predictors was calculated to detect influences of movement on the EEG signal. In the plank task, every recording segment was checked separately. The assumption of independent errors was tested using the Durbin-Watson statistic. Analyses were checked for collinearity and normality using the variance inflation factor (VIF) and P-P-plots.

## 4. Results

### 4. 1 Pre-registered analyses

#### Hypothesis 1 and 2

For the first recording segment (elevator), the repeated-measures 2 × 2 × 2 ANOVA with the within-subject factors validity (IAPS, plank), valence (neutral, negative), and electrode (F3/4, F7/8) and AIs as dependent variable, there was a main effect of electrode (F(1, 74) = 4.93, *p* = .03, ηp² = .06). Further, a significant interaction between validity, valence, and electrode was found (F(1, 74) = 5.84, *p* = .02, ηp² = .07). No post hoc test reached significance however after applying a Bonferroni correction (all *ps* > .12) (Figure 3A).

**Figure 3.**
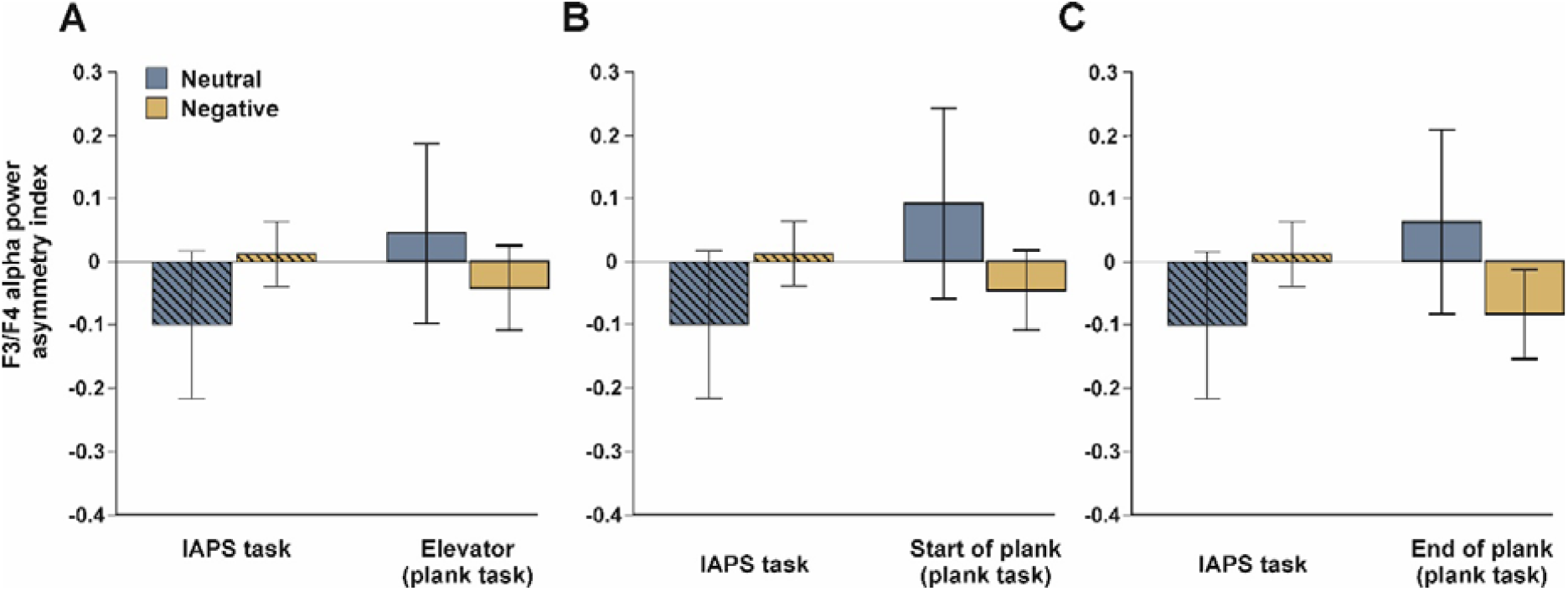
Asymmetry indices across both tasks for the neutral and negative conditions at the F3/F4 electrode sites. **A)** Comparison between IAPS results (hatched) and the ‘elevator’ segment from the plank task. **B)** Comparison between IAPS results and the ‘start of plank’ segment from the plank task. **C)** Comparison between IAPS results and the ‘end of plank’ segment from the plank task. Note that the IAPS results are identical for comparison as the IAPS only had a single recording segment. Error bars represent the standard error of the mean (SEM).

Analysis of the second recording segment (start of plank) revealed a main effect of electrode (F(1, 74) = 6.86, *p* = .01, ηp² = .09). A threefold interaction between validity, valence, and electrode was found (F(1, 74) = 4.11, *p* = .046, ηp² = .05). There was a trend toward significance in the more ecologically valid plank task on the electrode pair F3/4 (*p* = .06). Here, the AI decreased from the neutral (M = .09, SD = .66) to the negative condition (M = -.05, SD = .27) (Figure 3B).

The ANOVA for the ‘end of plank’ recording segment showed a main effect of electrode (F(1, 74) = 5.49, *p* = .02, ηp² = .07). Moreover, the interaction between validity, valence, and electrode reached significance (F(1, 74) = 4.2, *p* = .04, ηp² = .05), but post hoc tests did not survive Bonferroni correction. However, there was a trend toward significance that showed the same direction as for the second time point. Within the plank task, we found that AIs decreased from the neutral (M = .06, SD = .63) to the negative condition (M = -.08, SD = .31) on the F3/4 electrode (*p* = .051) suggesting greater left-hemispheric alpha power and thus stronger right-hemispheric activation in the negative condition (Figure 3C).

Specifically for the plank task, the 2 × 2 × 3 repeated-measures ANOVA with the within-subject factors valence (neutral, negative), electrode (F3/4, F7/8), and recording segment (‘elevator’, ‘start of plank’, ‘end of plank’) revealed a significant interaction between valence and recording segment (F(1.74, 128.76) = 7.84, *p* = .001, ηp² = .10). Bonferroni-corrected post hoc tests showed two significant differences within the negative condition pooled across the F3/4 and F7/8 electrode. The AI decreased significantly *(p* = .01) from ‘elevator’ (M = -.05, SD = 0.3) to ‘start of plank’ (M = -.08, SD = 0.3) *(p* = .012) and from ‘elevator’ to ‘end of plank’ (M = -.11, SD = 0.3) (*p* = .002). This indicates increasing involvement of the right hemisphere in the course of the negative plank condition (see Figure 4). The difference between ‘start of plank’ and ‘end of plank’ did not reach significance (*p* = .23). No significant differences between time points were found in the neutral condition (all *ps* > .11).

**Figure 4.**
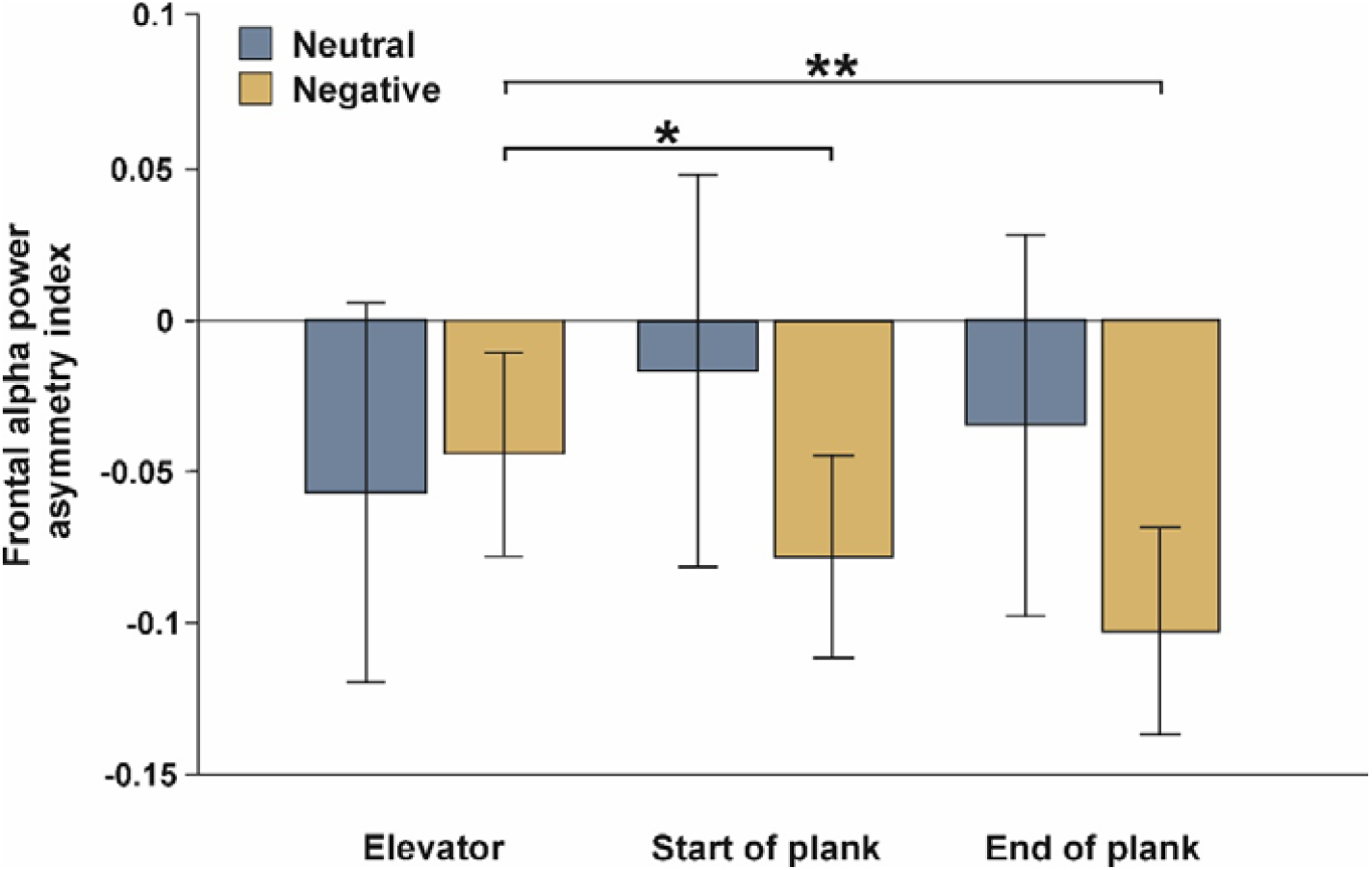
Temporal development of AIs during the plank task in the neutral and negative conditions. In the neutral condition, no changes in AIs were observed over time. In the negative condition however, significant decreases in AIs were observed in the ‘start of plank’ and ‘end of plank’ segment with respect to the elevator segment. Error bars represent SEM. * p < .05 and ** p < .01.

#### Hypothesis 3 and 4

AIs of the second (‘start of plank’: r = .23, *p* = .045) and third (‘end of plank’: r = .27, *p* = .02) recording segment of the negative plank condition were positively correlated with subjective fear ratings at these recording segments but only for the electrode pair F3/4 (see Figure 5). This implies that participants with greater subjective fear showed increasingly leftward frontal activity at these electrode sites. Correlations of AIs at the electrode pair F7/8 were not significant (all ps > .31). Concerning neutral conditions, none of the correlations reached significance (all *ps* > .15).

**Figure 5.**
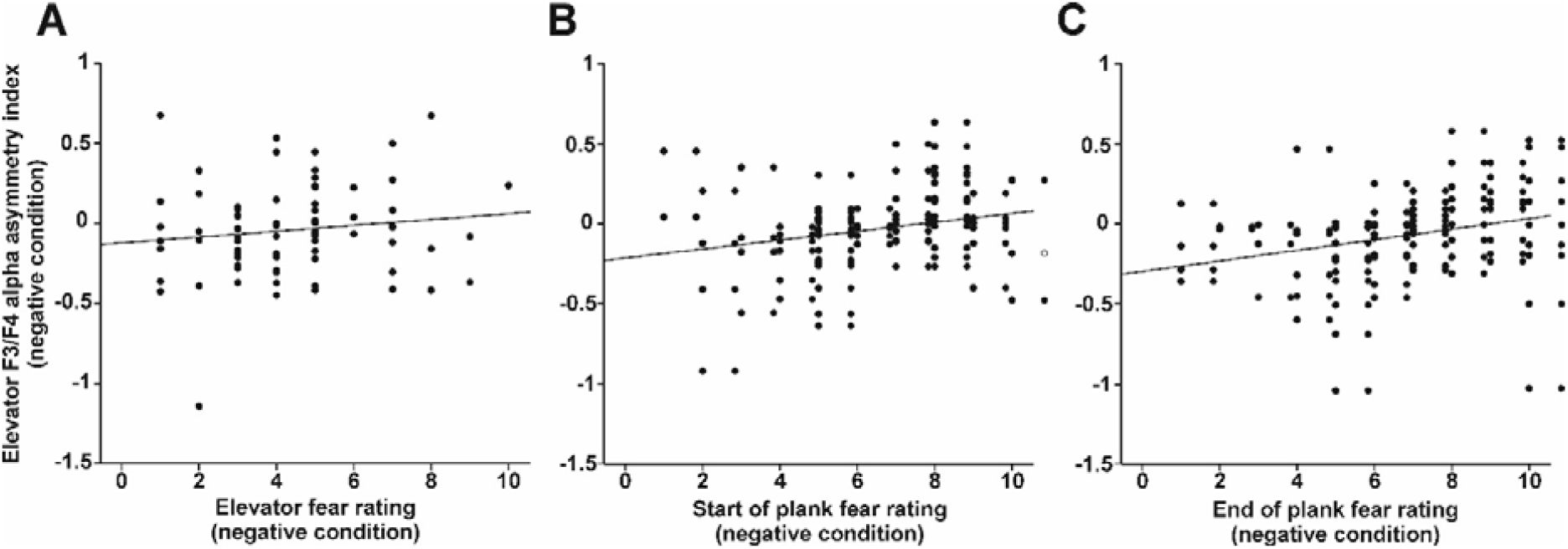
Pearson correlations between AIs at the F3/F4 electrode sites and subjective fear ratings across the recording segments. No significant association was detected between AIs and fear ratings during the **A)** ‘elevator’ segment. Significant positive correlations were observed for the **B)** ‘start of plank’ and **C)** ‘end of plank’ segment.

AIs in the negative IAPS condition were negatively correlated with the AIs in the negative plank condition on the electrode pair F3/4 (‘elevator’: r = -.25, *p* = .03; ‘start of plank’: r = -.23, *p* = .046; ‘end of plank’: r = -.28, *p* = .01) (see Figure 6). Thus, stronger right-hemispheric activation during VR was actually associated with stronger left-hemispheric activation during the IAPS task. There were no significant correlations between the two neutral conditions of the plank and the IAPS task (all *ps* > .12).

**Figure 6.**
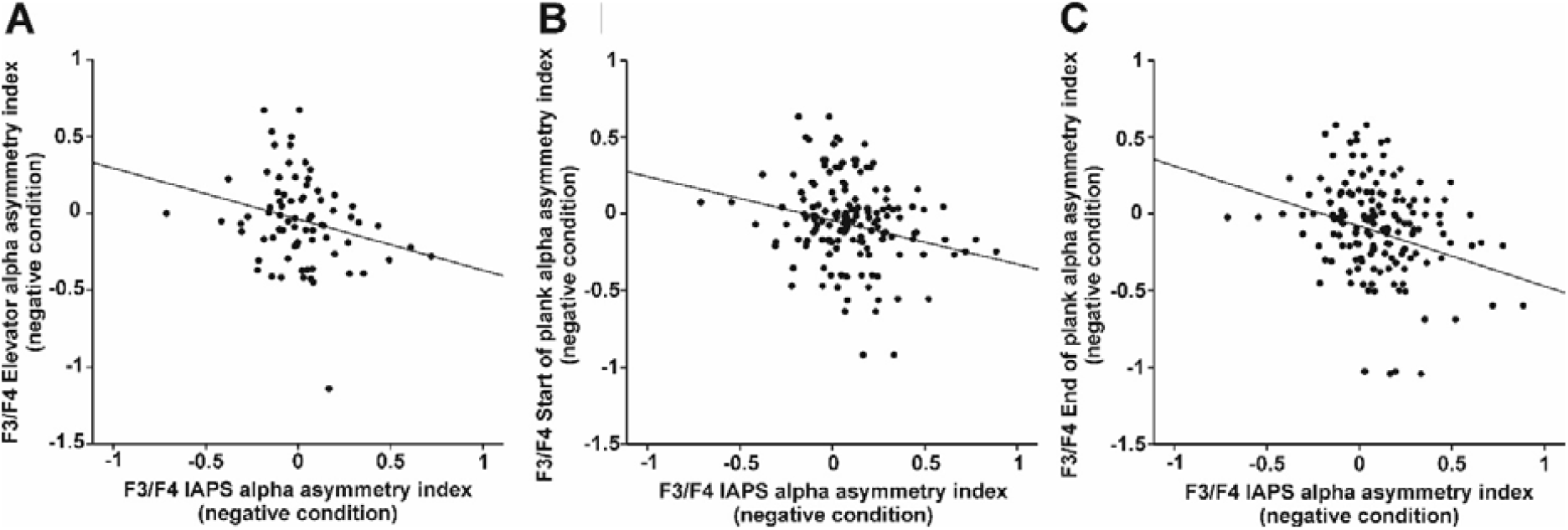
Pearson correlations between AIs at the F3/F4 electrode sites during the IAPS task and across the recording segments. Significant negative correlation coefficients could be detected during the **A)** ‘elevator’, the **B)** ‘start of plank’, and the **C)** ‘end of plank’ segment.

### 4. 2 Exploratory analyses

#### 4. 2. 1 Self-report data

Subjective fear ratings in the plank task were analysed with a 2 × 3 repeated-measures ANOVA with the within-subject factors valence (neutral, negative) and recording segment (‘elevator’, ‘start of plank’, ‘end of plank’) (for mean ratings see Table 1). A significant main effect of both valence (F(1, 73) = 286.97, *p* < .001, ηp² = .8) and recording segment (F(1.58, 115.41) = 72.93, *p* < .001, ηp² = .5) could be detected. There was a significant difference between the neutral (M = 1.6, SD = 0.77) and the negative plank condition with the latter showing higher self-reported fear (M = 5.64, SD = 2.12). Additionally, the factor valence interacted with the factor recording segment (F(2, 146) = 53.78, *p* < .001, ηp² = .42). Exclusively in the negative plank condition, the difference in fear was significant between all three recording segments. Thus, self-reported fear successively increased from ‘elevator’ (M = 4.41, SD = 2.1) to ‘start of plank’ (M = 6.03, SD = 2.28) (*p* < .001), from ‘start of plank’ to ‘end of plank’ (M = 6.47, SD = 2.47) (*p* = .02), and from ‘elevator’ to ‘end of plank’ (*p* < .001) (see Figure 7). At every recording segment, subjective fear ratings were greater in the negative than in the neutral plank condition (all *ps* < .001).

**Table 1.** Correlations of fear ratings in the plank conditions with pre- and post-questionnaires.

|  | plank (neutral) |  |  | plank (negative) |  |  |
| --- | --- | --- | --- | --- | --- | --- |
|  | elevator | start of plank | end of plank | elevator | start of plank | end of plank |
| acrophobia (ACRO/AVOI) | .25* | .2 | .27* | **.46\*\*\*** | **.49\*\*\*** | **.51\*\*\*** |
| avoidance (ACRO/AVOI) | .14 | .1 | .15 | .31** | **.38\*\*\*** | **.4\*\*\*** |
| presence (IPQ_G) | .05 | -.06 | -.03 | **.4\*\*\*** | **.39\*\*\*** | .34** |
| simulator sickness (SSQ) | .12 | .11 | .16 | .28* | .33** | .32** |

Note. \* = $p \leq .05$ , \*\* = $p \leq .01$ , \*\*\* = $p \leq .001$ . Bonferroni-corrected significant values ( $p = .0021$ ) are printed in bold.

**Figure 7.**
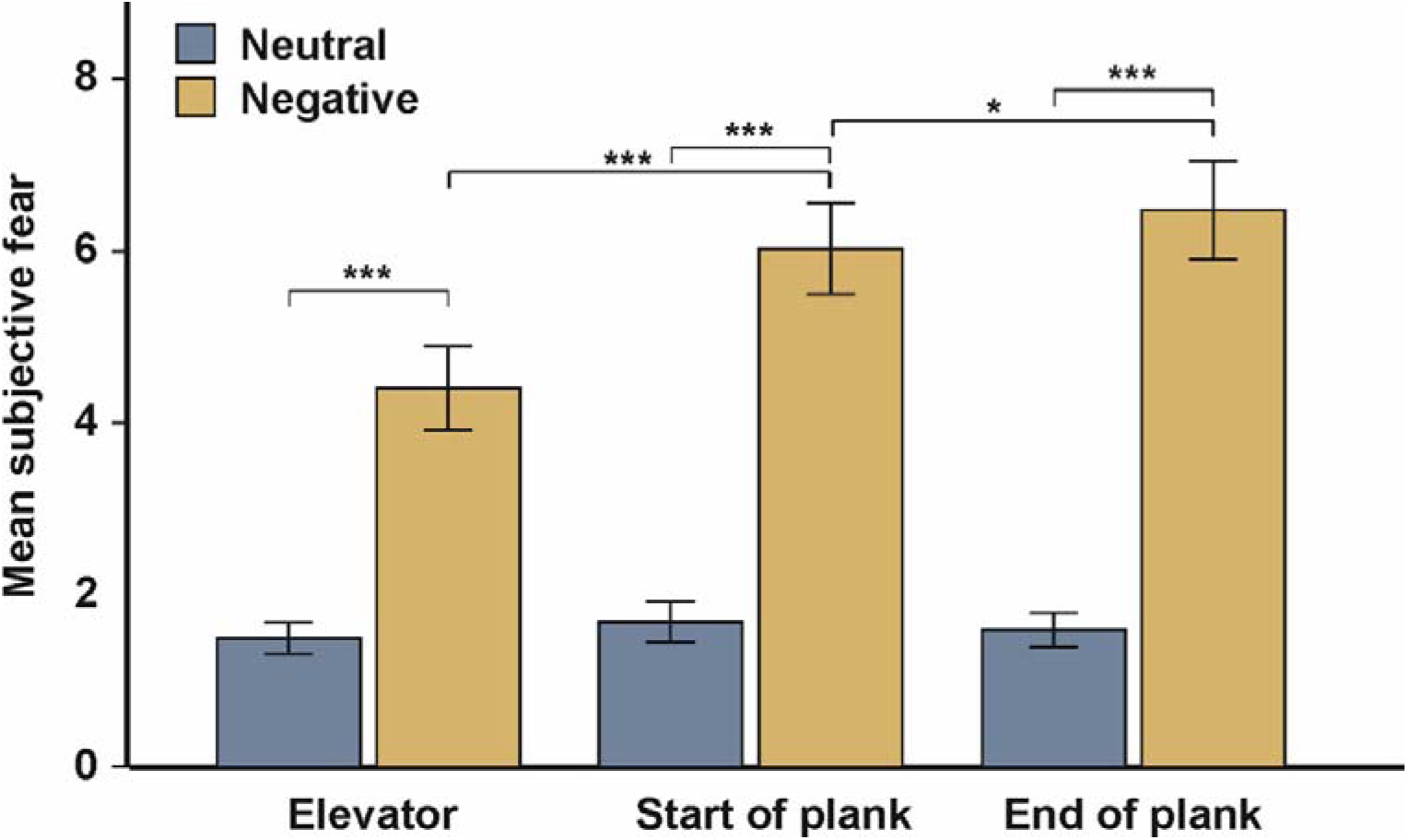
Subjective fear ratings for the neutral and negative condition across the recording segments. Subjective fear ratings did not change across the neutral condition and were at low levels overall. Fear ratings were substantially higher in the negative condition and successively increased over time. Error bars represent SEM. * p < .05, ** p < .01, and *** p < .001.

Correlations between fear ratings and the questionnaires were calculated. There was no significant correlation between the STAI-T score and self-reported fear regardless of condition (all *ps* > .33). The STAI-S was positively correlated with the ‘end of plank’ rating of both the neutral (r = .29, uncorrected *p* = .01) and the negative plank task (r = .33, uncorrected *p* < .01), and with the ‘start of plank’ rating in the negative condition only (r = .24, uncorrected *p* = .04). Thus, participants with higher state values of anxiety before the experiment later reported higher fear in the ratings especially in the negative condition. Specifically for fear of heights, the acrophobia and the avoidance subscale of the ACRO/AVOI questionnaire were positively correlated with all three ratings (see Table 1) of the negative plank condition. The acrophobia subscale was further significantly correlated with the ‘elevator’ and the ‘end of plank’ rating of the neutral condition. However, these correlations did not survive Bonferroni correction.

Fear ratings of the neutral plank task did not correlate with any of the presence ratings (all *ps* > .7). Correlations of the fear ratings in the negative plank task with corresponding presence ratings reached significance at every recording segment (‘elevator’: r = .47, *p* < .001; ‘start of plank’: r = .57, *p* < .001; ‘end of plank’: r = .55, *p* < .001). Thus, participants with higher self-reported presence also indicated higher fear. Overall presence ratings can be found in Supplementary table 1.

Correlations of fear ratings with post-experimental questionnaires yielded significant positive correlations between all subscales of the presence questionnaire (IPQ) and subjective presence ratings during the experiment, regardless of condition (see Table 1 and Table 2). Participants who stated a more pronounced feeling of presence during the experiment showed higher values in the IPQ which was filled out after the experiment. Significant correlations were found between two of the fear ratings in the negative plank condition (‘start of plank’ and ‘end of plank’) and the SSQ score (see Table 1). Higher subjective fear at these recording segments was accompanied by more pronounced physical symptoms (e.g., nausea, dizziness). After Bonferroni correction, these correlations were however not significant anymore.

**Table 2.** Correlations between presence ratings in the plank task and the subscales of the IPQ.

|  | plank (control) |  |  | plank (experimental) |  |  |
| --- | --- | --- | --- | --- | --- | --- |
|  | elevator | start of<br>plank | end of<br>plank | elevator | start of<br>plank | end of<br>plank |
| presence (IPQ_G) | **.42\*\*\*** | **.48\*\*\*** | **.5\*\*\*** | **.56\*\*\*** | **.58\*\*\*** | **.39\*\*\*** |
| presence (IPQ_SP) | **.4\*\*\*** | **.48\*\*\*** | **.52\*\*\*** | **.55\*\*\*** | **.51\*\*\*** | **.26\*** |
| presence (IPQ_INV) | .26* | .32** | .31** | **.38\*\*\*** | **.4\*\*\*** | .24* |
| presence (IPQ_REAL) | **.45\*\*\*** | **.5\*\*\*** | **.5\*\*\*** | **.57\*\*\*** | **.44\*\*\*** | **.36\*\*\*** |

*Note.* \* = $p \leq .05$ , \*\* = $p \leq .01$ , \*\*\* = $p \leq .001$ . Bonferroni-corrected significant values ( $p = .0021$ ) are printed in bold. G = general item, SP = spatial presence, INV = involvement, REAL = experienced realism. IPQ\_G is an additional general item and does not count as a subscale of the IPQ. IPQ\_SP measures the feeling of being in the simulation. IPQ\_INV inquires how focused and involved participants were in the VR. IPQ\_REAL represents how real the VR experience felt.

The repeated-measures 3 × 2 ANOVA of the PANAS questionnaires with the within-subject factors time (baseline, post-VR, post-IAPS) and affect (positive affect, negative affect) showed significant main effects of time (F(1.78, 131.86) = 53.63, *p* < .001, ηp² = .42) and affect (F(1, 74) = 391.77, *p* < .001, ηp² = .84). A significant interaction of time and affect was found (F(1.64, 121.26) = 85.84, *p* < .001, ηp² = .54). Post hoc Bonferroni correction for the main effect of time showed a significant increase of positive and negative affect from baseline (M = 1.86, SD = 0.27) to post-VR (M = 2.17, SD = 0.28) (*p* < .001) and a decrease from post-VR to post-IAPS (M = 1.82, SD = 0.36) (*p* < .001). Post hoc Bonferroni tests of affect revealed that participants generally reported higher positive affect (M = 2.59, SD = 0.46) than negative affect (M = 1.31, SD = 0.25) *(p* < .001). After post hoc Bonferroni correction of the time x affect interaction, results showed that positive affect increased from baseline (M = 2.6, SD = 0.5) to post-VR (M = 3.02, SD = 0.48) (*p* < .001) and then decreased at the post-IAPS timepoint (M = 2.15, SD = 0.64) (*p* < .001). The decrease of positive affect between baseline and post-IAPS also reached significance (*p* < .001). For negative affect, there was a successive increase from baseline (M = 1.13, SD = 0.2) to post-VR (M = 1.32, SD = 0.29) (*p* < .001) to post-IAPS rating (M = 1.49, SD = 0.52) (*p* = .01). The difference between baseline and post-IAPS was also significant (*p* < .001).

### 4. 2. 3 Movement signals

Additional multiple linear regressions were conducted to investigate the predictive value of movements (X-, Y-, and Z-axis) on the AIs. According to the Durbin-Watson statistic, residuals were independent. Approximate normality was checked via P-P-plots and histograms of the standardized residuals. Homoscedasticity and linearity were indicated based on the scatterplot of the residuals. According to the VIF, multicollinearity was not of concern (for the neutral condition: all VIF < 2; for the negative condition: all VIF < 2.77). In the neutral condition of the plank task, movements predicted AIs on the electrode pair F3/4 at two different recording segments (‘start of plank’: F(3,70) = 3.63, *p* = .02, R² = .13, ‘end of plank’: F(3, 71) = 2.83, *p* = .045, R² = .11) (see Table 5). After further analysis, only the X-axis significantly predicted AIs in the neutral plank task at these recording segments. No further significant influences of movement sensors could be detected (all *ps* > .1; see Table 6 for results in the negative plank condition). A significant influence of movements on the electrode pair F3/4 was found for the neutral IAPS task (F(3, 71) = 4.69, *p* = .001, R² = .165) (see Table 7). 16.5% of the variance in the AIs of the neutral IAPS task can be explained by the model. However, only the X-axis constituted a significant predictor (F3/4: *p* = .001). All regression models can be found in Supplementary tables 2 to 4.

## 5. Discussion

We aimed to investigate the neural correlates of fear processing both in an immersive and a 2D pictorial setting. In the immersive design, a virtual height paradigm was combined with a mobile EEG system to assess hemispheric asymmetries. These were compared to brain asymmetry patterns in response to two-dimensional emotional IAPS images to draw conclusions about the ecological validity of the established emotion induction method.

### 5. 1. Behavioral results during VR

We found clear indications that the fear induction of the VR paradigm worked exceptionally well using subjective ratings. Although research should not rely solely on subjective measures to evaluate emotion induction, fear ratings have generally been proven to be valuable ([62]; for a review, see [63]). Indeed, our results revealed substantially higher fear ratings in the negative VR condition compared to the neutral condition throughout all recording segments. Additionally, ratings showed an incremental increase in subjective fear over time. Thus, at least on a subjective level, participants experienced a considerable growth in fear when standing on the heightened plank. Notably, these patterns were not found in the neutral plank walk on the ground. Thus, the plank paradigm is a valid tool to induce fear not only between different conditions (negative vs. neutral) but also within conditions as standing within the safety of the elevator is perceived as less fearful compared to standing at the end of the plank.

### 5. 2. EEG results

#### 5. 2. 1. Confirmatory hypotheses

##### Hypothesis 1: Alpha asymmetries in the VR task

The first hypothesis targeted the efficacy of the VR paradigm in inducing fear reflected by altered hemispheric asymmetries in response to the virtual height simulation. Drawing on research on lateralization theories, higher left-hemispheric alpha power (reflecting greater right hemispheric activity) in the negative compared to the neutral VR condition was expected [64,65]. In the present study, the AI on electrode pair F3/4 indicated greater activation of the right hemisphere in the negative plank condition for the ‘start of plank’ and the ‘end of plank’ recording segment compared to the neutral condition but only on a trend level. This is in line with a VR study by RodrÍguez Ortega, Rey Solaz, and Alcañiz [66]. They reported that right-hemispheric alpha band activity decreased in response to VR-induced sad mood, supporting rightward dominance in negative emotion processing. The trends in the present experiment imply that there was a difference in brain activation between the neutral and the negative plank condition but only when standing on the plank when fear ratings were highest. Correspondingly, AIs in the negative plank condition successively shifted leftward indicating increasing involvement of the right hemisphere. However, this development was only significant between the first and the second and between the first and the third recording segment. Apparently, looking down the abyss from inside the elevator was not as powerful in evoking fear as standing on the plank. The time-dependent tendency toward right-hemispheric processing is well in line with the self-reported fear ratings as they were at the lowest in the elevator and successively increased on the plank.

##### Hypothesis 2: Alpha asymmetries in the IAPS task

The second hypothesis predicted more negative AIs in the negative IAPS condition compared to the neutral IAPS condition. Surprisingly, no significant differences in brain asymmetries were found between the neutral and the negative condition. Despite the widespread use of the IAPS, findings about emotion-related hemispheric asymmetries elicited by the images remain controversial. For positive emotions, Gable and Harmon-Jones [67] found leftward activation to appetitive two-dimensional stimuli. Alterations of brain asymmetry in multiple brain regions during IAPS exposure were also reported by Orgo, Bachmann, Lass, and Hinrikus [68]. At the same time, various studies have failed to detect asymmetric brain activation patterns during emotional picture presentation [69–71]. A possible explanation for this inconsistency in the literature and the null results in our data could be attributed to a lack of emotional impact of the IAPS images hindering naturalistic emotional brain responses.

##### Hypothesis 3: Relationship between fear ratings and alpha asymmetries in the plank task

The third hypothesis assumed a negative correlation between fear ratings and AIs in the negative plank condition. In the neutral condition, no significant correlations were revealed. For the ‘start of plank’ and ‘end of plank’ recording segment of the negative plank condition however, there were positive correlations between fear ratings and AIs indicating that greater fear was associated with greater left-hemispheric activation. The results were thus in the opposite direction from what we hypothesized. We can only speculate about the nature of this unexpected finding. One potential explanation for the enhanced left-hemispheric activation in participants with higher self-reported fear could be the application of emotion regulation strategies. Goodman, Rietschel, Lo, Costanzo, and Hatfield [72] found more pronounced leftward frontal asymmetry in particularly stressful situations. This activation pattern predicted better emotion regulation in participants and was accompanied by higher self-reported stress and anxiety. Similar findings were obtained by Berretz, Packheiser, Wolf, and Ocklenburg [73] who recorded EEG signals during the Trier Social Stress Test. If participants in the present study suppressed their emotions, applied cognitive reappraisal, or distracted themselves from standing on the plank, the EEG signal could be biased towards regulatory brain activation rather than reflecting a pure correlate of fear [74]. Thus, participants reporting higher self-reported fear could have applied emotion regulation strategies requiring greater left-hemispheric frontal involvement.

##### Hypothesis 4: Comparison of alpha asymmetries between tasks

The fourth hypothesis aimed at testing the ecological validity of the tasks. Ecological validity was postulated to differ significantly between the emotion induction methods used in this study, positive correlations were assumed between AIs of the negative IAPS condition and the negative plank condition. Higher correlations were expected between both neutral conditions. However, the data did not confirm this hypothesis as no correlations were detected between neutral conditions and we found low negative correlations for all three recording segments in the negative conditions. Since alpha oscillations show acceptable short-term reliability [75], it is unlikely that this result is due to spurious changes in oscillatory activity. It is possible that using a more naturalistic paradigm activates different brain networks, as patterns between the two tasks were only weakly related, in both negative and neutral settings.

#### 5. 2. 2. Movement signals

In this study, the impact of movements on the EEG signal was tested with multiple linear regressions. Results revealed significant effects implying a predictive value of movements on the EEG signal. However, these effects occurred only in the neutral conditions of both tasks and solely on electrode pair F3/4. A systematic impact of movement should have been evident consistently across conditions and recording segments. Similarly, several studies have demonstrated that movements do not significantly influence the EEG signal in mobile EEG or VR-based tasks [41,43,76]. Especially in asymmetry research, the use of a relative measure between hemispheres likely cancels out any residual movement artifacts. Thus, the selective findings between movements and the EEG signal in our study seem to be coincidental rather than systematic.

### 5. 3. Limitations and future directions

While in our study, we aimed to provide a highly naturalistic paradigm, one might reasonably argue that walking atop a plank on a skyscraper is not a realistic scenario. Even though fear and presence ratings were generally high, they could be further improved by implementing a variety of sensory cues. Previous studies [77,78] found a positive relationship between multimodal sensory stimulation and sense of presence in that a higher range in stimulated senses during VR resulted in a higher sense of presence. Especially auditory stimuli corresponding to the simulated situation seem to create “realness” [78]. In this experiment, auditory cues such as street noise or elevator noises could be implemented via headphones to create a more realistic surrounding. Additionally, a fan could give the impression of wind when leaving the elevator adding a somatosensory dimension. Thus, future studies could try to introduce a broader variation of sensory stimulation to enhance immersion into the VR.

In terms of measurements, there are possible improvements in the behavioral as well as in the physiological dimension. Since we speculated that emotional regulation might have played a role in the observed result pattern, a questionnaire assessing emotion regulation strategies (e.g., the German version of the Emotion Regulation Questionnaire [79]) would have been helpful to provide further support for this notion. On a physiological level, a complementary parameter recording peripheral responses to emotional material could also have been implemented. As discussed above, heart rate (or heart rate variability) and skin conductance constitute promising candidates for future research [80,81]. Finally, eye tracking could be considered as an additional measure to control for gaze direction. During the EEG recording on the plank, participants were instructed to freely explore their virtual surrounding to enable as much naturalistic behavior as possible. Therefore, control over the extent to which participants looked down the abyss was limited, and a certain amount of height avoidance was possible. This could be addressed by monitoring the participants’ gaze. Fortunately, this feature can easily be included in the VR system without requiring further equipment. This paradigm would also be highly suitable in the context of clinical laterality research to identify if, for example, phobic patients show altered lateralization patterns compared to healthy controls [82]. Furthermore, the reliable fear induction could also indicate that this paradigm is a suitable stress induction paradigm to study laterality patterns in more naturalistic settings.

All in all, the paradigm used in this study holds much potential for future laterality research. For example, as Mundorf and Ocklenburg [82] point out, there are many open questions on the role of lateralization in psychiatric and neurodevelopmental disorders. Thus, the field of clinical neuroscience could certainly benefit from a more naturalistic approach to the investigation of hemispheric asymmetries.

## 6. Conclusion

VR technology is already widespread in clinical research, especially in the treatment of anxiety disorders, and has proven to be effective on a behavioral level [24,83–86]. This study provides evidence that also the corresponding neurophysiological correlates of emotional processing are more reliably assessed in naturalistic settings that allow for feeling and corresponding action [87]. The combination of mobile EEG and VR could thus shed light on prevailing theories of emotional lateralization such as the right hemisphere and valence hypothesis in the future as this topic is still heavily debated.

## Data Accessibility

Data will be made fully available upon journal submission.

## Authors’ Contributions

Yasmin El Basbasse acquired, analyzed and interpreted the data and wrote the manuscript. Julian Packheiser supervised the data acquisition, analyzed the data, and wrote the manuscript. Jutta Peterburs interpreted the data and conceived the study design. Onur Güntürkün provided laboratory space and equipment and interpreted the data. Christopher Maymon programmed the VR simulation and interpreted the data. Gina Grimshaw interpreted the data, conceived the study design, and supervised the project. Sebastian Ocklenburg interpreted the data, conceived the study design, and supervised the project. All authors critically reviewed the manuscript.

## References

[1] Hughlings-Jackson J. 1878 On affections of speech from disease of the brain. Brain 1, 304–330. (http://dx.doi.org/10.1093/brain/1.3.304).

[2] Heilman KM, Scholes R, Watson RT. 1975 Auditory affective agnosia. Disturbed comprehension of affective speech. Journal of neurology, neurosurgery, and psychiatry 38, 69– 72. (http://dx.doi.org/10.1136/jnnp.38.1.69).

[3] DeKosky ST, Heilman KM, Bowers D, Valenstein E. 1980 Recognition and discrimination of emotional faces and pictures. Brain and Language 9, 206–214. (http://dx.doi.org/10.1016/0093-934X(80)90141-8).

[4] Palomero-Gallagher N, Amunts K. 2022 A short review on emotion processing: a lateralized network of neuronal networks. Brain Struct Funct 227, 673–684. (http://dx.doi.org/10.1007/s00429-021-02331-7).

[5] Ley RG, Bryden MP. 1979 Hemispheric differences in processing emotions and faces. Brain and Language 7, 127–138. (http://dx.doi.org/10.1016/0093-934X(79)90010-5).

[6] Schwartz GE, Davidson RJ, Maer F. 1975 Right hemisphere lateralization for emotion in the human brain: interactions with cognition. Science 190, 286–288. (http://dx.doi.org/10.1126/science.1179210).

[7] Sackeim HA, Gur RC, Saucy MC. 1978 Emotions are expressed more intensely on the left side of the face. Science 202, 434–436. (http://dx.doi.org/10.1126/science.705335).

[8] Roether CL, Omlor L, Giese MA. 2008 Lateral asymmetry of bodily emotion expression. Current Biology 18, R329–30. (http://dx.doi.org/10.1016/j.cub.2008.02.044).

[9] Gainotti G. 2012 Unconscious processing of emotions and the right hemisphere. Neuropsychologia 50, 205–218. (http://dx.doi.org/10.1016/j.neuropsychologia.2011.12.005).

[10] Costafreda SG, Brammer MJ, David AS, Fu CHY. 2008 Predictors of amygdala activation during the processing of emotional stimuli: a meta-analysis of 385 PET and fMRI studies. Brain Research Reviews 58, 57–70. (http://dx.doi.org/10.1016/j.brainresrev.2007.10.012).

[11] Dimond SJ, Farrington L. 1977 Emotional response to films shown to the right or left hemisphere of the brain measured by heart rate. Acta Psychologica 41, 255–260. (http://dx.doi.org/10.1016/0001-6918(77)90020-8).

[12] Tomarken AJ, Davidson RJ, Henriques JB. 1990 Resting frontal brain asymmetry predicts affective responses to films. Journal of Personality and Social Psychology 59, 791–801. (http://dx.doi.org/10.1037/0022-3514.59.4.791).

[13] Terzian H. 1964 Behavioural and EEG effects of intracarotid sodium amytal injection. Acta neurochir 12, 230–239. (http://dx.doi.org/10.1007/BF01402095).

[14] Prete G, Laeng B, Fabri M, Foschi N, Tommasi L. 2015 Right hemisphere or valence hypothesis, or bothã The processing of hybrid faces in the intact and callosotomized brain. Neuropsychologia 68, 94–106. (http://dx.doi.org/10.1016/j.neuropsychologia.2015.01.002).

[15] Ahern GL, Schwartz GE. 1979 Differential lateralization for positive versus negative emotion. Neuropsychologia 17, 693–698. (http://dx.doi.org/10.1016/0028-3932(79)90045-9).

[16] Demaree HA, Everhart DE, Youngstrom EA, Harrison DW. 2005 Brain lateralization of emotional processing: historical roots and a future incorporating “dominance". Behavioral and Cognitive Neuroscience Reviews 4, 3–20. (http://dx.doi.org/10.1177/1534582305276837).

[17] Lang PJ, Bradley MM, Cuthbert BN. 2008 International affective picture system (IAPS): Affective ratings of pictures and instruction manual. Technical Report A-8. University of Florida, Gainesville, FL.

[18] Courtney CG, Dawson ME, Schell AM, Iyer A, Parsons TD. 2010 Better than the real thing: eliciting fear with moving and static computer-generated stimuli. International journal of psychophysiology 78, 107–114. (http://dx.doi.org/10.1016/j.ijpsycho.2010.06.028).

[19] Lang PJ, Greenwald MK, Bradley MM, Hamm AO. 1993 Looking at pictures: affective, facial, visceral, and behavioral reactions. Psychophysiology 30, 261–273. (http://dx.doi.org/10.1111/j.1469-8986.1993.tb03352.x).

[20] Bradley MM, Cuthbert BN, Lang PJ. 1996 Picture media and emotion: effects of a sustained affective context. Psychophysiology 33, 662–670. (http://dx.doi.org/10.1111/j.1469-8986.1996.tb02362.x).

[21] Detenber BH, Simons RF, Bennett GG. 1998 Roll ‘em!: The effects of picture motion on emotional responses. Journal of Broadcasting & Electronic Media 42, 113–127. (http://dx.doi.org/10.1080/08838159809364437).

[22] Horvat M, Kukolja D, Ivanec D. 2015 Comparing affective responses to standardized pictures and videos: A study report. Proceedings of the 38th International Convention, IEEE, 1394– 1398.

[23] Fernández C, Pascual JC, Soler J, Elices M, Portella MJ, Fernández-Abascal E. 2012 Physiological responses induced by emotion-eliciting films. Appl Psychophysiol Biofeedback 37, 73–79. (http://dx.doi.org/10.1007/s10484-012-9180-7).

[24] Diemer J, Lohkamp N, Mühlberger A, Zwanzger P. 2016 Fear and physiological arousal during a virtual height challenge--effects in patients with acrophobia and healthy controls. Journal of anxiety disorders 37, 30–39. (http://dx.doi.org/10.1016/j.janxdis.2015.10.007).

[25] Klinger E, Bouchard S, Légeron P, Roy S, Lauer F, Chemin I, Nugues P. 2005 Virtual reality therapy versus cognitive behavior therapy for social phobia: a preliminary controlled study. Cyberpsychology & behavior : the impact of the Internet, multimedia and virtual reality on behavior and society 8, 76–88. (http://dx.doi.org/10.1089/cpb.2005.8.76).

[26] Salehi E, Mehrabi M, Fatehi F, Salehi A. 2020 Virtual Reality Therapy for Social Phobia: A Scoping Review. In Digital Personalized Health and Medicine, pp. 713–717: IOS Press.

[27] Clus D, Larsen ME, Lemey C, Berrouiguet S. 2018 The Use of Virtual Reality in Patients with Eating Disorders: Systematic Review. Journal of medical Internet research 20, e7898. (http://dx.doi.org/10.2196/jmir.7898).

[28] Abulrub A-HG, Attridge AN, Williams MA. 2011 Virtual reality in engineering education: The future of creative learning. In 2011 IEEE Global Engineering Education Conference (EDUCON): IEEE.

[29] McCloy R, Stone R. 2001 Science, medicine, and the future. Virtual reality in surgery. BMJ 323, 912–915. (http://dx.doi.org/10.1136/bmj.323.7318.912).

[30] Kavanagh S, Luxton-Reilly A, Wuensche B, Plimmer B. 2017 A systematic review of Virtual Reality in education. Themes in Science and Technology Education 10, 85–119. See https://www.learntechlib.org/p/182115/.

[31] Liang M, Starrett MJ, Ekstrom AD. 2018 Dissociation of frontal-midline delta-theta and posterior alpha oscillations: A mobile EEG study. Psychophysiology 55, e13090. (http://dx.doi.org/10.1111/psyp.13090).

[32] Diersch N, Wolbers T. 2019 The potential of virtual reality for spatial navigation research across the adult lifespan. J Exp Biol 222. (http://dx.doi.org/10.1242/jeb.187252).

[33] Bischof WF, Boulanger P. 2003 Spatial navigation in virtual reality environments: an EEG analysis. Cyberpsychology & behavior 6, 487–495. (http://dx.doi.org/10.1089/109493103769710514).

[34] Baños RM, Botella C, Alcañiz M, Liaño V, Guerrero B, Rey B. 2004 Immersion and emotion: their impact on the sense of presence. Cyberpsychology & behavior 7, 734–741. (http://dx.doi.org/10.1089/cpb.2004.7.734).

[35] MarÍn-Morales J, Higuera-Trujillo JL, Greco A, Guixeres J, Llinares C, Scilingo EP, Alcañiz M, Valenza G. 2018 Affective computing in virtual reality: emotion recognition from brain and heartbeat dynamics using wearable sensors. Sci Rep 8, 13657. (http://dx.doi.org/10.1038/s41598-018-32063-4).

[36] MarÍn-Morales J, Higuera-Trujillo JL, Greco A, Guixeres J, Llinares C, Gentili C, Scilingo EP, Alcañiz M, Valenza G. 2019 Real vs. immersive-virtual emotional experience: Analysis of psycho-physiological patterns in a free exploration of an art museum. PLOS ONE 14, e0223881. (http://dx.doi.org/10.1371/journal.pone.0223881).

[37] Barke A, Stahl J, Kröner-Herwig B. 2012 Identifying a subset of fear-evoking pictures from the IAPS on the basis of dimensional and categorical ratings for a German sample. Journal of Behavior Therapy and Experimental Psychiatry 43, 565–572. (http://dx.doi.org/10.1016/j.jbtep.2011.07.006).

[38] Horvat M, Dobrinic M, Novosel M, Jercic P. 2018 Assessing emotional responses induced in virtual reality using a consumer EEG headset: A preliminary report. In 2018 41st International Convention on Information and Communication Technology, Electronics and Microelectronics (MIPRO). May 21-25, 2018, Opatija, Croatia : proceedings. Piscataway, NJ: IEEE.

[39] Hofmann SM, Klotzsche F, Mariola A, Nikulin VV, Villringer A, Gaebler M. 2020 Decoding subjective emotional arousal from EEG during an immersive Virtual Reality experience. bioRxiv, 2020.10.24.353722. (http://dx.doi.org/10.1101/2020.10.24.353722).

[40] Debener S, Minow F, Emkes R, Gandras K, de Vos M. 2012 How about taking a low-cost, small, and wireless EEG for a walkã Psychophysiology 49, 1617–1621. (http://dx.doi.org/10.1111/j.1469-8986.2012.01471.x).

[41] Packheiser J, Schmitz J, Pan Y, El Basbasse Y, Friedrich P, Güntürkün O, Ocklenburg S. 2020 Using Mobile EEG to Investigate Alpha and Beta Asymmetries During Hand and Foot Use. Front. Neurosci. 14, 109. (http://dx.doi.org/10.3389/fnins.2020.00109).

[42] Tremmel C, Herff C, Krusienski DJ. 2019 EEG Movement Artifact Suppression in Interactive Virtual Reality. In 41st Annual International Conference of the IEEE Engineering in Medicine and Biology Society (EMBC): IEEE.

[43] Wang W-E, Ho RLM, Gatto B, van Der Veen SM, Underation MK, Thomas JS, Antony AB, Coombes SA. 2020 A Novel Method to Understand Neural Oscillations During Full-Body Reaching: A Combined EEG and 3D Virtual Reality Study. IEEE transactions on neural systems and rehabilitation engineering 28, 3074–3082. (http://dx.doi.org/10.1109/tnsre.2020.3039829).

[44] Koller-Schlaud K, Querbach J, Behr J, Ströhle A, Rentzsch J. 2020 Test-Retest Reliability of Frontal and Parietal Alpha Asymmetry during Presentation of Emotional Face Stimuli in Healthy Subjects. NPS 79, 428–436. (http://dx.doi.org/10.1159/000505783).

[45] Ocklenburg S, Güntürkün O. 2017 The Lateralized Brain. The Neuroscience and Evolution of Hemispheric Asymmetries. Saint Louis: Elsevier Science.

[46] Zhao G, Zhang Y, Ge Y. 2018 Frontal EEG Asymmetry and Middle Line Power Difference in Discrete Emotions. Front. Behav. Neurosci. 12, 225. (http://dx.doi.org/10.3389/fnbeh.2018.00225).

[47] Balconi M, Mazza G. 2010 Lateralisation effect in comprehension of emotional facial expression: a comparison between EEG alpha band power and behavioural inhibition (BIS) and activation (BAS) systems. Laterality 15, 361–384. (http://dx.doi.org/10.1080/13576500902886056).

[48] Beck AT, Ward C, Mendelsohn M, Mock J, Erbaugh JJAGP. 1961 Beck depression inventory (BDI). Archives of General Psychiatry 4, 561–571.

[49] Coan JA, Allen JJB. 2004 Frontal EEG asymmetry as a moderator and mediator of emotion. Biological Psychology 67, 7–49. (http://dx.doi.org/10.1016/j.biopsycho.2004.03.002).

[50] Kong X-Z et al. 2020 Mapping brain asymmetry in health and disease through the ENIGMA consortium. Hum Brain Mapp. (http://dx.doi.org/10.1002/hbm.25033).

[51] Baker BL, Cohen DC, Saunders JT. 1973 Self-directed desensitization for acrophobia. Behaviour Research and Therapy 11, 79–89. (http://dx.doi.org/10.1016/0005-7967(73)90071-5).

[52] Hüweler R, Kandil FI, Alpers GW, Gerlach AL. 2009 The impact of visual flow stimulation on anxiety, dizziness, and body sway in individuals with and without fear of heights. Behaviour Research and Therapy 47, 345–352. (http://dx.doi.org/10.1016/j.brat.2009.01.011).

[53] Lang PJ, Bradley MM. 2007 The International Affective Picture System (IAPS) in the study of emotion and attention. In Handbook of Emotion Elicitation and Assessment (eds JA Coan, JJB Allen), pp. 29–46. New York: Oxford University Press.

[54] SteamVR [Online]: Valve Corporation. Available: https://store.steampowered.com/steamvr.

[55] Spielberger CD, Gorsuch RL, Lushene PR, Vagg PR, Jacobs AG. 1983 Manual for the State-Trait Anxiety Inventory (Form Y). Palo Alto: Consulting Psychologists Press.

[56] Krohne HW, Egloff B, Kohlmann C-W, Tausch A. 1996 PsycTESTS Dataset: American Psychological Association (APA).

[57] Watson D, Clark LA, Tellegen A. 1988 Development and validation of brief measures of positive and negative affect: The PANAS scales. Journal of Personality and Social Psychology 54, 1063–1070. (http://dx.doi.org/10.1037/0022-3514.54.6.1063).

[58] Kennedy RS, Lane NE, Berbaum KS, Lilienthal MG. 1993 Simulator Sickness Questionnaire: An Enhanced Method for Quantifying Simulator Sickness. The International Journal of Aviation Psychology 3, 203–220. (http://dx.doi.org/10.1207/s15327108ijap0303_3).

[59] Regenbrecht HT, Schubert TW, Friedmann F. 1998 Measuring the Sense of Presence and its Relations to Fear of Heights in Virtual Environments. International Journal of Human-Computer Interaction 10, 233–249. (http://dx.doi.org/10.1207/s15327590ijhc1003_2).

[60] Jasper H. 1958 Report of the committee on methods of clinical examination in electroencephalography. Electroencephalography and Clinical Neurophysiology 10, 370–375. (http://dx.doi.org/10.1016/0013-4694(58)90053-1).

[61] Reznik SJ, Allen JJB. 2018 Frontal asymmetry as a mediator and moderator of emotion: An updated review. Psychophysiology 55, e12965. (http://dx.doi.org/10.1111/psyp.12965).

[62] McLean CP, Hope DA. 2010 Subjective anxiety and behavioral avoidance: Gender, gender role, and perceived confirmability of self-report. Journal of anxiety disorders 24, 494–502. (http://dx.doi.org/10.1016/j.janxdis.2010.03.006).

[63] Hersen M. 1973 Self-assessment of fear. Behavior therapy 4, 241–257.

[64] Coan JA, Allen JJB. 2003 The state and trait nature of frontal EEG asymmetry in emotion. In The Asymmetrical Brain (eds K Hugdahl, RJ Davidson), pp. 565–615: The MIT Press.

[65] Palmiero M, Piccardi L. 2017 Frontal EEG Asymmetry of Mood: A Mini-Review. Front. Behav. Neurosci. 11, 224. (http://dx.doi.org/10.3389/fnbeh.2017.00224).

[66] RodrÍguez Ortega A, Rey Solaz B, Alcañiz M. 2013 Evaluating Virtual Reality Mood Induction Procedures with Portable EEG Devices. Annual Review of Cybertherapy and Telemedicine 11, 131–135. See http://hdl.handle.net/10251/78725.

[67] Gable P, Harmon-Jones E. 2008 Relative left frontal activation to appetitive stimuli: considering the role of individual differences. Psychophysiology 45, 275–278. (http://dx.doi.org/10.1111/j.1469-8986.2007.00627.x).

[68] Orgo L, Bachmann M, Lass J, Hinrikus H. 2015 Effect of negative and positive emotions on EEG spectral asymmetry. Annual International Conference of the IEEE Engineering in Medicine and Biology Society., 8107–8110. (http://dx.doi.org/10.1109/EMBC.2015.7320275).

[69] Winkler I, Jager M, Mihajlovic V, Tsoneva T. 2010 Frontal EEG asymmetry based classification of emotional valence using common spatial patterns. World Academy of Science, Engineering and Technology 45, 373–378. (http://dx.doi.org/10.5281/zenodo.1061729).

[70] Murphy FC, Nimmo-Smith I, Lawrence AD. 2003 Functional neuroanatomy of emotions: a meta-analysis. Cognitive, Affective, & Behavioral Neuroscience 3, 207–233. (http://dx.doi.org/10.3758/cabn.3.3.207).

[71] Huster RJ, Stevens S, Gerlach AL, Rist F. 2009 A spectralanalytic approach to emotional responses evoked through picture presentation. International journal of psychophysiology 72, 212–216. (http://dx.doi.org/10.1016/j.ijpsycho.2008.12.009).

[72] Goodman RN, Rietschel JC, Lo L-C, Costanzo ME, Hatfield BD. 2013 Stress, emotion regulation and cognitive performance: the predictive contributions of trait and state relative frontal EEG alpha asymmetry. International journal of psychophysiology 87, 115–123. (http://dx.doi.org/10.1016/j.ijpsycho.2012.09.008).

[73] Berretz G, Packheiser J, Wolf OT, Ocklenburg S. 2022 Acute stress increases left hemispheric activity measured via changes in frontal alpha asymmetries. iScience 25, 103841. (http://dx.doi.org/10.1016/j.isci.2022.103841).

[74] Turnbull OH, Salas CE. 2021 The Neuropsychology of Emotion and Emotion Regulation: The Role of Laterality and Hierarchy. Brain Sciences 11, 1075. (http://dx.doi.org/10.3390/brainsci11081075).

[75] Mathewson KJ, Hashemi A, Sheng B, Sekuler AB, Bennett PJ, Schmidt LA. 2015 Regional electroencephalogram (EEG) alpha power and asymmetry in older adults: a study of short-term test-retest reliability. Front. Aging Neurosci. 7, 177. (http://dx.doi.org/10.3389/fnagi.2015.00177).

[76] Packheiser J, Berretz G, Rook N, Bahr C, Schockenhoff L, Güntürkün O, Ocklenburg S. 2021 Investigating real-life emotions in romantic couples: a mobile EEG study. Sci Rep 11, 1142. (http://dx.doi.org/10.1038/s41598-020-80590-w).

[77] Cooper N, Milella F, Pinto C, Cant I, White M, Meyer G. 2018 The effects of substitute multisensory feedback on task performance and the sense of presence in a virtual reality environment. PLOS ONE 13, e0191846. (http://dx.doi.org/10.1371/journal.pone.0191846).

[78] Honegger F, Feng Y, Rauterberg M. 2021 Multimodality for Passive Experience: Effects of Visual, Auditory, Vibration and Draught Stimuli on Sense of Presence. jucs 27, 582–608. (http://dx.doi.org/10.3897/jucs.68384).

[79] Abler B, Kessler H. 2009 Emotion Regulation Questionnaire – Eine deutschsprachige Fassung des ERQ von Gross und John. Diagnostica 55, 144–152. (http://dx.doi.org/10.1026/0012-1924.55.3.144).

[80] Peterson SM, Furuichi E, Ferris DP. 2018 Effects of virtual reality high heights exposure during beam-walking on physiological stress and cognitive loading. PLOS ONE 13, e0200306. (http://dx.doi.org/10.1371/journal.pone.0200306).

[81] Fadeev KA, Smirnov AS, Zhigalova OP, Bazhina PS, Tumialis AV, Golokhvast KS. 2020 Too Real to Be Virtual: Autonomic and EEG Responses to Extreme Stress Scenarios in Virtual Reality. Behavioural Neurology 2020, 5758038. (http://dx.doi.org/10.1155/2020/5758038).

[82] Mundorf A, Ocklenburg S. 2021 The clinical neuroscience of lateralization. London, New York: Routledge Taylor & Francis Group.

[83] Emmelkamp P, Krijn M, Hulsbosch A, de Vries S, Schuemie M, van der Mast C. 2002 Virtual reality treatment versus exposure in vivo: a comparative evaluation in acrophobia. Behaviour Research and Therapy 40, 509–516. (http://dx.doi.org/10.1016/s0005-7967(01)00023-7).

[84] Emmelkamp P, Meyerbröker K. 2021 Virtual Reality Therapy in Mental Health. Annual review of clinical psychology 17, 495–519. (http://dx.doi.org/10.1146/annurev-clinpsy-081219-115923).

[85] Hodges LF, Kooper R, Meyer TC, Rothbaum BO, Opdyke D, de Graaff JJ, Williford JS, North MM. 1995 Virtual environments for treating the fear of heights. Computer 28, 27–34. (http://dx.doi.org/10.1109/2.391038).

[86] Opriş D, Pintea S, GarcÍa-Palacios A, Botella C, Szamosközi Ş, David D. 2012 Virtual reality exposure therapy in anxiety disorders: a quantitative meta-analysis. Depression and anxiety 29, 85–93. (http://dx.doi.org/10.1002/da.20910).

[87] Ocklenburg S, Berretz G, Packheiser J, Friedrich P. 2021 Laterality 2020: entering the next decade. Laterality 26, 265–297. (http://dx.doi.org/10.1080/1357650X.2020.1804396).

