## Supplementary information for "Walk the Plank! Using mobile EEG to investigate emotional lateralization of immersive fear in virtual reality"

**Supplementary Table 1****.** Mean fear and presence ratings for the three recording segments of the neutral and the negative plank condition.

|  | fear | | presence | |
| --- | --- | --- | --- | --- |
|  | neutral | negative | neutral | negative |
| elevator | 1.5  (0.78) | 4.41  (2.1) | 6.14  (1.95) | 6.76  (1.88) |
| start of plank | 1.69  (1) | 6.03  (2.28) | 6.61  (1.92) | 7.53  (1.8) |
| end of plank | 1.59  (0.86) | 6.47  (2.47) | 6.64  (2.02) | 7.61  (1.8) |

*Note*. Standard deviations are indicated in parenthesis.

**Supplementary Table 2.** Multiple regression of AIs (F3/4, F7/8) on movements (X-, Y-, and Z-axis) at the three recording segments in the neutral plank condition.

|  | elevator | | | start of plank | | | end of plank | | |
| --- | --- | --- | --- | --- | --- | --- | --- | --- | --- |
|  | Beta (ß) | *p* | *t* | Beta (ß) | *p* | *t* | Beta (ß) | *p* | *t* |
| F3/4 |  | | | | | | | | |
| intercept |  | .71 | .38 |  | .046* | 2.03 |  | .72 | .36 |
| X-axis | .27 | .03* | 2.26 | .33 | .01** | 2.92 | .33 | .01** | 2.76 |
| Y-axis | -.02 | .9 | -.12 | -.22 | .08* | -1.76 | .-.01 | .95 | -.06 |
| Z-axis | .15 | .19 | 1.32 | .2 | .12 | 1.6 | .1 | .54 | .61 |
| F7/8 |  | | | | | | | | |
| intercept |  | .29 | -1.07 |  | .4 | .84 |  | .04* | 2.12 |
| X-axis | -.09 | .48 | -.72 | .07 | .54 | .62 | .13 | .3 | 1.05 |
| Y-axis | .12 | .34 | .97 | -.11 | .43 | -.8 | -.33 | .04* | -2.06 |
| Z-axis | .14 | .26 | 1.14 | .22 | .1 | 1.65 | .37 | .02* | 2.33 |

*Note.* * = *p* ≤ .05, ** = *p* ≤ .01.

**Supplementary Table 3.** Multiple regression of AIs (F3/4, F7/8) on movements (X-, Y-, and Z-axis) at the three recording segments in the negative plank condition.

|  | elevator | | | start of plank | | | end of plank | | |
| --- | --- | --- | --- | --- | --- | --- | --- | --- | --- |
|  | Beta (ß) | *p* | *t* | Beta (ß) | *p* | *t* | Beta (ß) | *p* | *t* |
| F3/4 |  | | | | | | | | |
| intercept |  | .67 | .43 |  | .41 | -.84 |  | .43 | .79 |
| X-axis | .1 | .43 | .8 | .12 | .3 | 1.05 | .18 | .12 | 1.59 |
| Y-axis | -.08 | .69 | -.41 | .15 | .39 | .86 | -.12 | .46 | -.74 |
| Z-axis | -.53 | .6 | -.53 | .2 | .23 | 1.21 | -.05 | .74 | -.33 |
| F7/8 |  | | | | | | | | |
| intercept |  | .64 | -.47 |  | .38 | -.88 |  | .79 | .27 |
| X-axis | .08 | .52 | .64 | .1 | .38 | .88 | .18 | .12 | 1.57 |
| Y-axis | .1 | .62 | .51 | .14 | .4 | .84 | -.04 | .78 | -.28 |
| Z-axis | .12 | .51 | .66 | .28 | .1 | 1.68 | .2 | .2 | 1.31 |

**Supplementary table 4.** Multiple regression of AIs (F3/4, F7/8) on movements (X-, Y-, and Z-axis) in the IAPS task.

|  | neutral | | | negative | | |
| --- | --- | --- | --- | --- | --- | --- |
|  | Beta  (ß) | *p* | *t* | Beta  (ß) | *p* | *t* |
| F3/4 |  |  |  |  |  |  |
| intercept |  | .11 | 1.63 |  | .81 | .25 |
| X-axis | .44 | .001*** | 3.47 | -.06 | .65 | -.46 |
| Y-axis | -.19 | .13 | -.1.53 | -.04 | .79 | -.27 |
| Z-axis | -.1 | .35 | -.94 | -.06 | .63 | -.49 |
| F7/8 |  |  |  |  |  |  |
| intercept |  | .53 | .63 |  | .54 | -.61 |
| X-axis | .29 | .04* | 2.13 | .11 | .42 | .81 |
| Y-axis | -.08 | .57 | -.57 | .08 | .54 | .62 |
| Z-axis | .07 | .54 | .61 | .13 | .29 | 1.07 |

*Note.* * = *p* ≤ .05, *** = *p* ≤ .001
